# Pharmacokinetic profile of the synthetic mu-opioid receptor agonist [Lys7]Dermorphin-IRDye®800CW and its feasibility as a biomarker for opioid use disorder

**DOI:** 10.1101/2023.06.19.545627

**Authors:** Kimberley S. Samkoe, Rendall R. Strawbridge, Mark R. Spaller, Alexandre A. Pletnev, Dennis J. Wirth, Brook K. Byrd, Boyu Meng, J. Scott Sottosanti, Scott C. Davis, Jonathan T. Elliott

**Author notes:** **Corresponding Author:** Prof. Jonathan T. Elliott, Department of Orthopaedics, Dartmouth-Hitchcock Medical Center, 1 Medical Center Drive, Lebanon, NH 03756, Email: Jonathan Elliott.

## Abstract

**Background:** Opioid use disorder (OUD) affects more than 14 million Americans and poses a high risk of relapse, overdose, and death. Current treatments are not tailored to individual needs and do not monitor the effectiveness of the medication. We propose a novel method to measure the occupancy of mu opioid receptors (MOR), which are key targets for opioid pharmacotherapy, in peripheral tissues with high MOR density. We developed a fluorescent peptide agonist that binds to MOR and can be detected by non-invasive point-of-care techniques. We present *in vitro* and *in vivo* results that demonstrate the feasibility and potential of this method to assess MOR availability and treatment efficacy in OUD patients.

**Methods:** A new fluorescent-labeled synthetic peptide agonist [Lys7]Dermorphin-IRDye800CW, called DRM-800, was synthesized and characterized *in vitro* to evaluate binding and internalization. Wildtype and MOR knock-out mice were used to quantify plasma kinetics and, using a cyromacrotome, fluorescence images were acquired post-mortem on whole-body sections 150 um apart. These volumes were used to compare *in vivo* enhancement of MOR-rich structures.

**Results:** In vitro assays and microscope visualization of DRM-800 showed high MOR-affinity and rapid, robust internalization. Plasma half-life following intravenous injection in mice was 8-12 minutes. Specific binding by tissue structures of interest, measured by the ratio of relative fluorescent units in wild-type vs. MOR knockout mice showed high binding in dorsal root ganglia, spiral ganglia and trigeminal ganglion, as well as in the small and large intestine.

**Conclusions:** The pharmacokinetics and distribution, binding kinetics and rapid internalization suggests that MOR-specific fluorescence enhancement corresponding to opioid rich structures could serve as a potential biomarker in opioid use disorder.

## Introduction

In 2021, approximately 80,000 individuals died in the United States from opioid overdose, and the estimated prevalence of opioid use disorder (OUD) is now greater than 14 million (1,2). Unlike other substance use disorders, the misused drug will continue to play a legitimate and necessary role in the alleviation of acute pain and the long-term treatment of chronic pain in about 50 million adults in the US (3,4). However, it is now clear that the overuse and underappreciation of the addictive potential of opioids has resulted in what is commonly referred to as the ‘opioid epidemic’; for many who developed OUD as a result, their life will likely be one of cycling in and out of pharmacotherapy, relapse and overdose risk (5).

While maintenance medications are first-line standard-of-care therapy to prevent relapse and illicit drug use in individuals with OUD (6), there is no ‘one-size-fits-all’ approach, and currently no evidence to predict which patient will do best with which medication (7). The reason for this is thought to be individual differences at the pharmacological level—mu opioid receptor (MOR) expression is subject-variable and dynamic during exposure, treatment and withdrawal. For example, when OUD patients are treated with buprenorphine, a partial agonist for MOR that also blocks it from being activated by other drugs, the availability of MOR is correlated highly to opioid withdrawal symptoms at both the group and individual-subject levels, but the *slope and intercept* vary widely between individuals (6). Pharmacotherapy for OUD would be more effective if an individual’s MOR availability could directly inform dosing or tapering down efforts (8). To this end, we have developed a biomarker that has the potential to be used as a peripheral MOR sensor in benchtop animal studies as well as in a future first-in-human Phase 0 investigation.

Molecularly targeted fluorophores have emerged in cancer imaging as an effective way to leverage the onco-specific overexpression of cell-surface receptors to produce imaging contrast, enhancing the intraoperative visibility of malignant cells (9). The principle is based on the idea that fluorophores can be conjugated to small protein fragments or antibodies without substantial change in binding affinity (10), and then can be injected intravenously where specific uptake by tissue will directly correspond to affinity and concentration of available receptors (11,12). For example, in a study by Tichauer *et al.*, cetuximab, a clinical-grade monoclonal antibody against epidermal growth factor receptor (EGFR), was conjugated to IRDye^®^800CW and injected in the lymph nodes of mice with metastatic breast cancer of a cell line known to overexpress EGFR (13). As few as 200 cells (corresponding to approximately 3×10^7^ receptors) were detectable using this method. In an effort to develop *in vivo* biomarker assays to improve evidence-based pharmacotherapy in opioid use disorder, a similar approach is presented in this paper. Unlike in the case of cancer, where overexpression of receptors drives fluorescence signal, we focus on receptor occupancy or availability as the main corollate of available receptor concentration.

To assess the feasibility of this approach, we demonstrate *in vitro* and *in vivo* results characterizing the pharmacokinetic and pharmacodynamic properties of the proposed biomarker [Lys7]Dermorphin-IRDye^®^800CW, referred to as “DRM-800”. As a corollary, we present *in silico* analysis of the relationship between tissue fluorescence behaviour measured in an MOR-rich structure such as the spiral ganglia or dorsal root ganglia, and the occupancy of MORs by a competitive agonist or antagonist such as the clinically important methadone, buprenorphine, naltrexone or naloxone.

## Materials and Methods

### Preparation of DER-800

Preparation of the unlabeled peptide was performed using standard Fmoc-based solid-phase peptide synthesis (SPPS) methods as follows: Rink Amide AM resin (0.141 g, 0.1 mmol, 0.71 mmol/g loading) was swollen in DMF for 30 min, then drained. This was followed by initial Fmoc deprotection (20% piperidine/DMF, shaken for 1 min; drained; washed with DMF; repeated once). After DMF wash (shaken for 15 sec; drained; repeated twice), sequential coupling of the remaining residues began. This involved adding the appropriate Fmoc amino acid (0.5 mmol) precombined with HCTU (0.5 mmol) in DMF (1.5 mL) to the resin. After mixing for 20 seconds, DIEA (0.17 mL, 1 mmol) was added and the mixture was shaken for 3 min. After DMF wash (shaken for 15 sec; drained, repeated twice) the previous piperidine deprotection conditions were used. These steps were repeated for each added amino acid. After final Fmoc deprotection, the resin was sequentially washed with DMF and DCM (twice each). Peptide removal from the resin with global deprotection was accomplished with the addition of resin cleavage solution (5x resin volume, TFA/triisopropylsilane/thioanisole/anisole, volume ratio 92:4:2:2) and shaking for 1 h at r.t. The TFA solution was collected, mixed with cold diethyl ether, and the precipitated crude YdAFGYPKC-NH_2_ was separated by centrifugation, dissolved in water and lyophilized. LCMS of the crude peptide LCMS (Phenomenex Kinetex 2.6u XB-C18 100A column, 100×2.10 mm; acetonitrile gradient 0 to 70% in H_2_O-formic acid (0.1%) in 15 min, flow rate 0.2 mL/min): retention time 9.0 min, m/z calculated 947, found 474 (M+2H^+^), 948 (M+H^+^); 946 (M-H^+^).

Labeling with IRDye^®^800CW maleimide was performed using previously described methods (Ye et al., 2011). Briefly, to a solution of the crude YdAFGYPKC-NH_2_ (10 mg) in PBS buffer (pH 7.2-7.4, 10 mL) was added IRDye^®^800CW (18 mg), and the mixture was incubated for 2.5 h at r.t. The resulting solution was diluted with 20 mL of 0.1% TFA in water and purified by HPLC (Phenomenex Jupiter Proteo 4u AXIA 90A column, 250×21.20 mm; methanol gradient 50 to 100% in H_2_O-TFA (0.1%) in 20 min, flow rate 15 mL/min, collecting the fractions eluting between 9 and 13 min). The product-containing fractions were combined and lyophilized to afford 2 mg of IRDye^®^800CW-labelled YdAFGYPKC-NH_2_. LCMS (Phenomenex Kinetex 2.6u XB-C18 100A column, 100×2.10 mm; acetonitrile gradient 0 to 70% in H_2_O-formic acid (0.1%) in 15 min, flow rate 0.2 mL/min): retention time 12.5 min, m/z calculated 2072, found 1035 (M-2H^+^).

### In vitro studies

A Chinese hamster ovary (CHO) cell line with a stable mu-opioid receptor expression level (B_max_) of approximately 5.7 ± 3.0 pmol/mg protein measured by the manufacturer using [3H]-DAMGO (ES-542-C OP3 ValiScreen, PerkinElmer, Waltham, MA) were obtained and cultured for a series of binding assays. The cells were cultured in F-12 HAM’s medium supplemented with 1 mM L-glutamine, 10% (v/v) fetal bovine serum and 400 μg/ml G418, at 37⁰C. Two competitive assays were performed to determine the binding kinetics of DRM-800 *in vitro*. First, cells were incubated with 50 nM of DRM-800 and concentrations of naloxone in concentrations ranging from 100 pM to 5 μM in DPBS with Ca^2+^ and Mg^2+^ simultaneously. After 30 min incubation, the cells were washed with three times and a final 100 μL of DPBS with Ca^2+^ and Mg^2+^ added then imaged on the LI-COR Odyssey. Specific binding was calculated from the percent fluorescence signal compared to controls (DRM-800 with no naloxone, and no naloxone or DRM-800). A second assay was performed exactly as the first, except that the cells were first incubated with DRM-800 for 30 minutes followed by naloxone for 30 minutes, to determine the relative effect of internalization on receptor availability.

Cells were visualized using a Zeiss LSM 800 inverted microscope with a Generation III 750 nm laser light source (Lumencor, Inc., Beaverton, OR) and a Zeiss Plan-Apochromat 20×/0.8 M27 objective lens. The filters used for imaging DRM-800 were 740 nm for excitation, 770 nm dichroic and 780 nm longpass for emission (Filter Set 49037, Chroma Technology Corp., Bellows Falls, VT). Acquisition of micrographs was accomplished using a digital CMOS Camera (C114400-22CU, Hamamatsu Photonics K.K.) with a 2×2 pixel binning. Images were processed in ImageJ FIJI, windowed and leveled identically across samples, and then adjusted step-wise until cells are visible, with each image being shown for transparency.

### In Vivo Imaging

All animal experiments were approved by Dartmouth’s Institutional Animal Care and Use Committee (IACUC) under Protocol No. 00002070. Three different strains of mice were obtained from Jackson Laboratories: B6.129S2 (wild-type), B6.129S2-Oprm1^tm1Kff^IJ (MOR knockout), and B6.129S2-Oprm1^tm4Kff^IJ (mCherry knockin). Mice were assigned to one of four groups: wildtype, wildtype + naloxone, MOR-knockout, and mCherry knockin. To characterize the plasma concentration following injection, the direct carotid artery imaging method was used that has been previously described (14). Briefly, following induction of a surgical plane of anesthesia using 1.5-2% isoflurane, a sharp dissection of the neck was performed, and superficial tissue was removed to allow access to the underlying vasculature. Blunt dissection was performed using forceps to expose one of the common carotid arteries, separating it from the adjacent connective tissue. A piece of black polyvinyl cloth was placed behind the artery, so that only fluorescence from the artery is detected and covered in clear, plastic film to maintain hydration. The animal was placed in the LI-COR Pearl planar fluorescence imaging system, supine, and while anesthesia was maintained, was restrained using medical tape and the body covered in black polyvinyl to reduce fluorescence contribution from the body. A 1 nmol equimolar coinjection of DRM-800 and IRDye^®^700DX was performed directly into the tail vein, and the imaging stage was immediately moved into the scanner and closed (within 30 s of the injection). Fluorescence images were acquired for both 700 nm and 800 nm channels every 30-35 s for 1 hour at a resolution of 84 microns. Mice were immediately removed from the imaging device and frozen at −20 ℃ in OCT gel for cyro-macrotome imaging.

### Cyro-Macrotome Imaging

The cryo-imaging system (15,16) consisted of a modified whole animal cryo-macrotome (CM3600 Leica Biosystems, Wetzlar, Germany) fitted with a custom epi-illumination hyperspectral imaging system to acquire white light and fluorescence images during sectioning. The detection system consisted of two optical channels separated by a 750 nm dichroic mirror, one for visible and one for NIR imaging, each of which was fitted with a scientific CMOS camera (PCO.Edge 4.2 s, Kelheim, Germany). The optical path in the visible channel also contained a high-speed motorized filter wheel (HS1024, FLI, Lima, NY) and a liquid crystal tunable filter (VariSpect LCTF, PerkinElmer, Waltham, MA) for hyperspectral imaging. In this study, white light reflectance data was collected by illuminating with a white light LED (Mightex, Toronto, ON) and acquiring images between 420 nm and 720 nm in 10-nm increments using the LCTF. DRM-800 was imaged by illuminating the specimen with a beam-expanded 760 nm laser (CrystaLaser, LC, Reno, NV) and filtering with a 780 nm longpass filter (Chroma, Bellows Falls, VT) on the detection channel. Automated imaging of a full specimen volume was accomplished by; (1) acquiring white light reflectance images, (2) acquiring a DRM-800 fluorescence image, (3) sectioning, and (4) repeating on the newly-exposed surface. Imaging an entire mouse at 150 µm thick sections generally required between 150 and 170 slices.

For each image, dark light background subtraction was applied followed by flat field, radial distortion, exposure time, and camera registration corrections. White light images sampled within the visible range (420 nm - 720 nm) were spectrally remixed to recreate colored RGB images using the CIE 1931 Standard Observer function. After these basic processing steps, both RGB and fluorescence image stacks could then be reconstructed into high-resolution, whole-body, three-dimensional volumes with 100 um-scale resolution or better.

### Image Analysis

To extract plasma concentration curves from the Pearl imaging data, an in-house developed ICA-augmented algorithm was used. First, pixels associated with the carotid vessel region of interest are selected using the *imfreehand* tool in MATLAB (MathWorks, Natick, MA). Next, because the Pearl is a closed-box planar imaging system, as opposed to a raster-scanned or structured-light based system, fluorescence from across the mouse can influence the signal. Although this is mitigated somewhat by the placement of a black cloth under the vessel, and draping of the mouse, any extraneous fluorescence signal is unmixed from the dominant plasma-curve signal by applying fastICA, an independent component analysis algorithm. This was performed for all animals and a normalized group average curve was obtained. The procedure was repeated for both DRM-800 and IRDye^®^700DX.

To determine the whole body biodistribution of DRM-800 following intravenous injection, using the volumetric data (both fluorescence and RGB) obtained by the macrocyrotome method described above structures of interest were identified in the RGB volumes and fluorescence in the corresponding regions were quantified. The main structures of interest were brain, dorsal root ganglia (DRG), left and right spiral ganglia (SG), lungs, liver, stomach, small and large intestine, kidney, bladder, muscle and skin. Enhancement in the wildtype and mCherry-knockin mice were compared with the knockout mice.

### In Silico Experiments

A forward model was constructed using a modified Lammertsma two-tissue three-compartment model (17), where an additional tissue/compartment is added to represent internalized receptor/ligand that is still fluorescently active. The result is a plasma compartment (*C*_p_) input, and tissue compartments designated ‘free-space’ (*C*_f_), ‘bound-space (*C*_b_)’ and ‘internalized-space (*C*_i_)’. Rate constants *K*_1_ and *k*_2_ govern the influx and efflux from *C_p_* to *C_f_*, *k_3_*and *k_4_* govern the influx and efflux from *C_f_*to *C_b_*, and *k_int_* governs the influx from *C_b_* into *C_i_*. Binding potential of available receptors, *BP_ND_*, was defined as *k_3_/k_4_*, and the relationship between MOR availability and buprenorphine tablet dose was based on reported relationships from PET imaging (18). Forward data was generated for typical values *K_1_-k_4_* and *k_int_*, with *k_3_* being modified according to MOR availability, 10% gaussian white noise was added and Logan Graphical Analysis (19) was performed on the data to obtain a slope for each simulation.

### Statistics

All statistics in this study were carried out using SPSS or MATLAB. Biexponential curve fitting to recover plasma curve rate constants was performed using the *cftool* in MATLAB and used a Trust-Region algorithm with a *TolFun* setting of 1.0 x 10^−6^ and a *DiffMinChange* and *DiffMaxChange* of 1.0 x 10^−8^ and 0.1, respectively. *In vitro* assays were run in triplicate and data-points represent mean ± SD. Specific binding was calculated using ImageJ Fiji (version 1.51n) as mean pixel value within the circular ROI. and GraphPad Prism (version 8.0, San Diego, CA) was used to fit a competitive binding model. Shaded regions behind the plasma curves in Figure 4 represent the standard deviation, whereas values appearing as pairs in parenthesis separated by a comma represent the 95% confidence interval of the mean.

## Results

### Competitive binding with naloxone shows strong MOR affinity and rapid internalization

An *in vitro* competitive binding assay was performed using DRM-800 as the labeled agent and either unlabeled [Lys7]Dermorphin or naloxone as the competitive ligand. Figure 2A shows the results of the competitive assay when 2 nM of DRM-800 is incubated with [Lys7]Dermorphin across a range of concentrations. The IC-50 of [Lys7]Dermorphin was 40 nM. Figure 2B reports the results of a competitive assay using naloxone as the competitor. First, CHO-K1 cells were simultaneously incubated with 50 nM of DRM-800 and various concentrations of naloxone (0-1000 nM): the IC_50_ was 3.5 nM. However, when cells were incubated with 50 nM of DRM-800 only, for 30 minutes, and then sequentially incubated with varying concentrations of naloxone (0-1000 nM) for 30 minutes, no competitive inhibition was observed and the specific binding of DRM-800 was not influenced by naloxone concentration.

**Fig 1.**
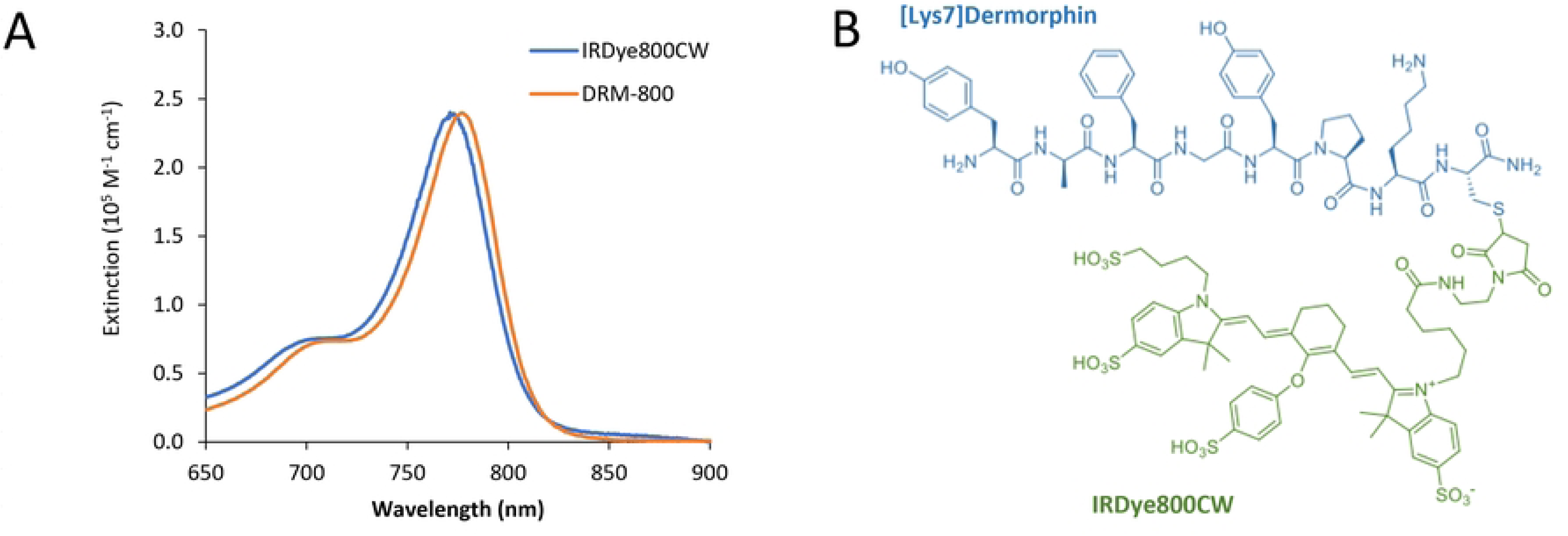
Molecular and optical properties. (A) The absorption spectra for commercial IRDye800CW and DRM-800 (B) The chemical structure of DRM-800, showing the maleimide link between the cystine terminal amino acid and IRDye800CW. (B) The absorption and emission spectra of IRDye^®^800CW before and after conjugation with [Lys7]Dermorphin.

**Fig. 2.**
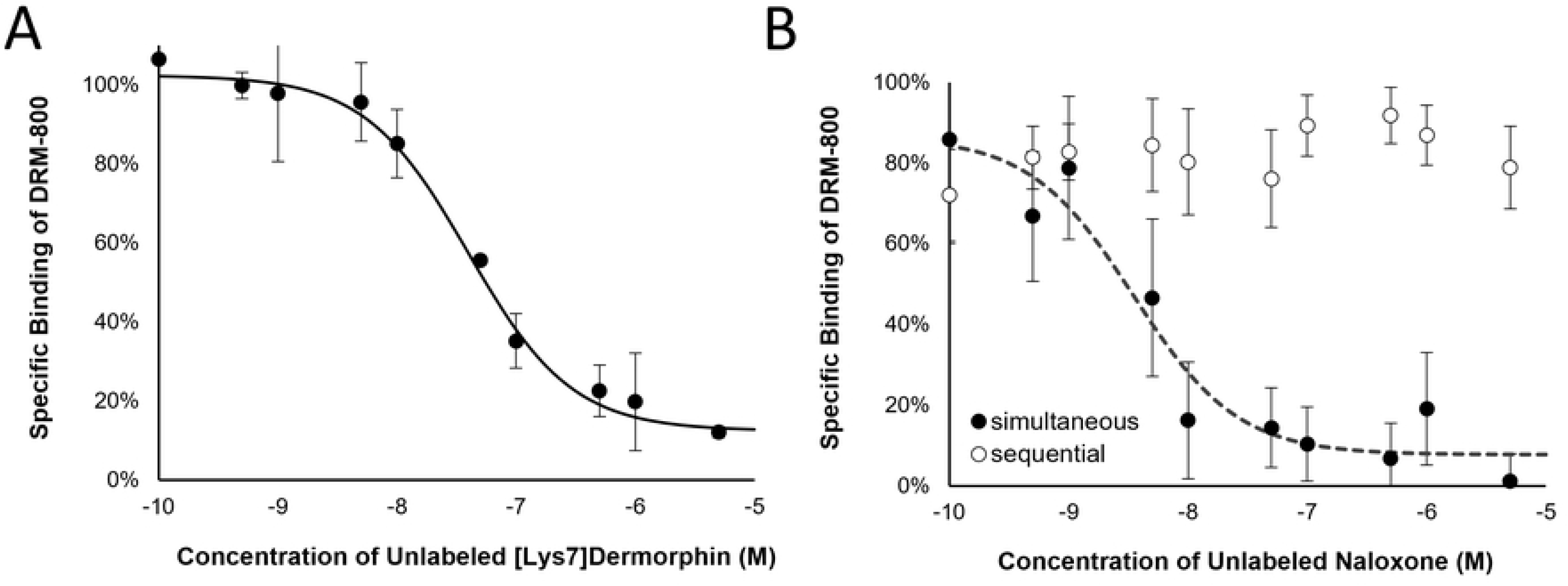
Competitive binding. Results of two competitive binding assays. (A) Cells were incubated with 2 nM concentrations of DRM-800 and simultaneously with [Lys7]Dermorphin across a range of concentrations. (B) Cells were either simultaneously incubated with DRM-800 at 50 nM and a range of naloxone concentrations, or they were sequentially incubated with DRM-800 being administered 30 minutes before naloxone for another 30 minutes.

Wide field fluorescence microscopy (Fig. 3) was used to study the binding and internalization of DRM-800 to stably expressed MORs in the membrane of transfected CHO-K1 cells. At a concentration of 20 and 200 times the reported K_d_ of [Lys7]Dermorphin (20) (2 nM and 25 nM) and 35°C, plasma membrane fluorescence as well as fluorescent puncta of high intensity in the perinuclear region of the cytoplasm were observable after 30 min of incubation. At low temperatures (4C) with the addition of sucrose (0.35 M) to inhibit internalization (21,22), fluorescence was distinctly observed at the plasma membrane and intracellular puncta was greatly reduced.

**Fig. 3.**
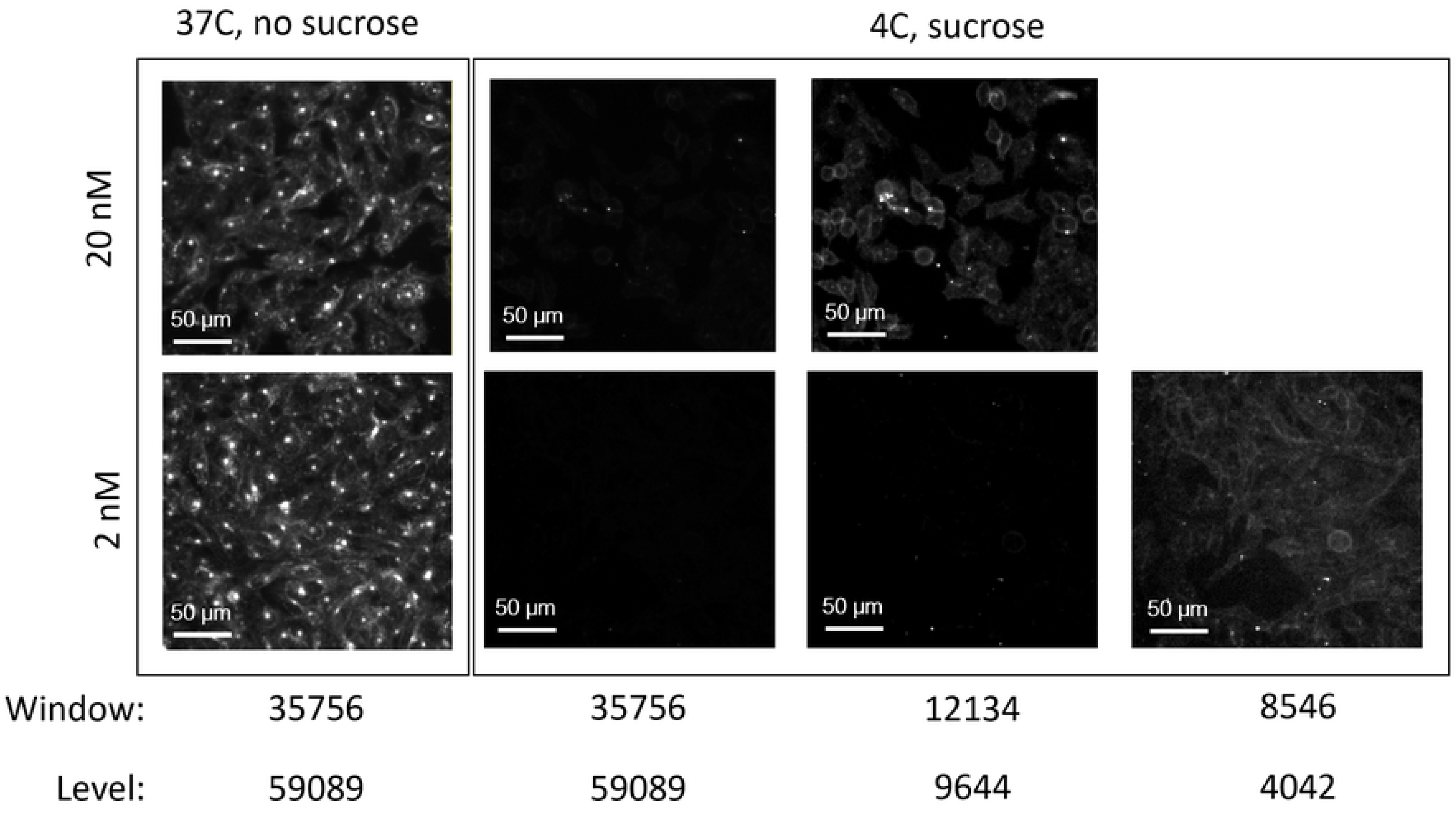
Fluorescence images. Fluorescence micrographs of CHO-K1 cells expressing MOR, taken at 20x for two different concentrations of DRM-800 and with or without sucrose to block internalization. Images are windowed and leveled according to the values specified in the bottom two lines.

### Plasma pharmacokinetics were obtained following intravenous administration of DRM-800

Plasma curves obtained using the direct carotid imaging method were subsequently fit with a biexponential model, *y = a exp(-bx) + c exp(-dx)* to obtain distribution rate, *b*, and elimination rate, *d*. The recovered values for DRM-800 were 0.211 (0.197 - 0.224) min^−1^ and 0.0097 (0.0093 - 0.0103) min^−1^ for *b* and *d*, respectively in the wildtype mice; whereas in the MOR-knockout mice *b* and *d* were 0.11 (0.08-0.14) min^−1^ and 0.0155 (0.0102 - 0.0202) min^−1^, respectively. For IRD700 the rate constants did not differ between groups, being 0.42 (0.36 - 0.47) min^−1^ and 0.054 (0.051 - 0.057) min^−1^ for *b* and *d*, respectively. The plasma half-life (time in which the concentration of dye is 50% of initial) for DRM-800 was 7.95 min and 12.6 min for wildtype and knockout mice, respectively. However, the biexponential half-lives (half-lives for each compartment) were also calculated. The distribution (α) and elimination (ꞵ) half-lives were 3.3 (3.1-3.5) min and 70.8 (67.2 - 74.8) min, respectively, for wildtype mice and 6.2 (4.9 - 8.6) min and 45 (33–68) min, respectively, for knockout mice. Figure 4 shows the mean plasma curves for DRM-800 and IRDye700DX for each group of mice.

**Fig. 4.**
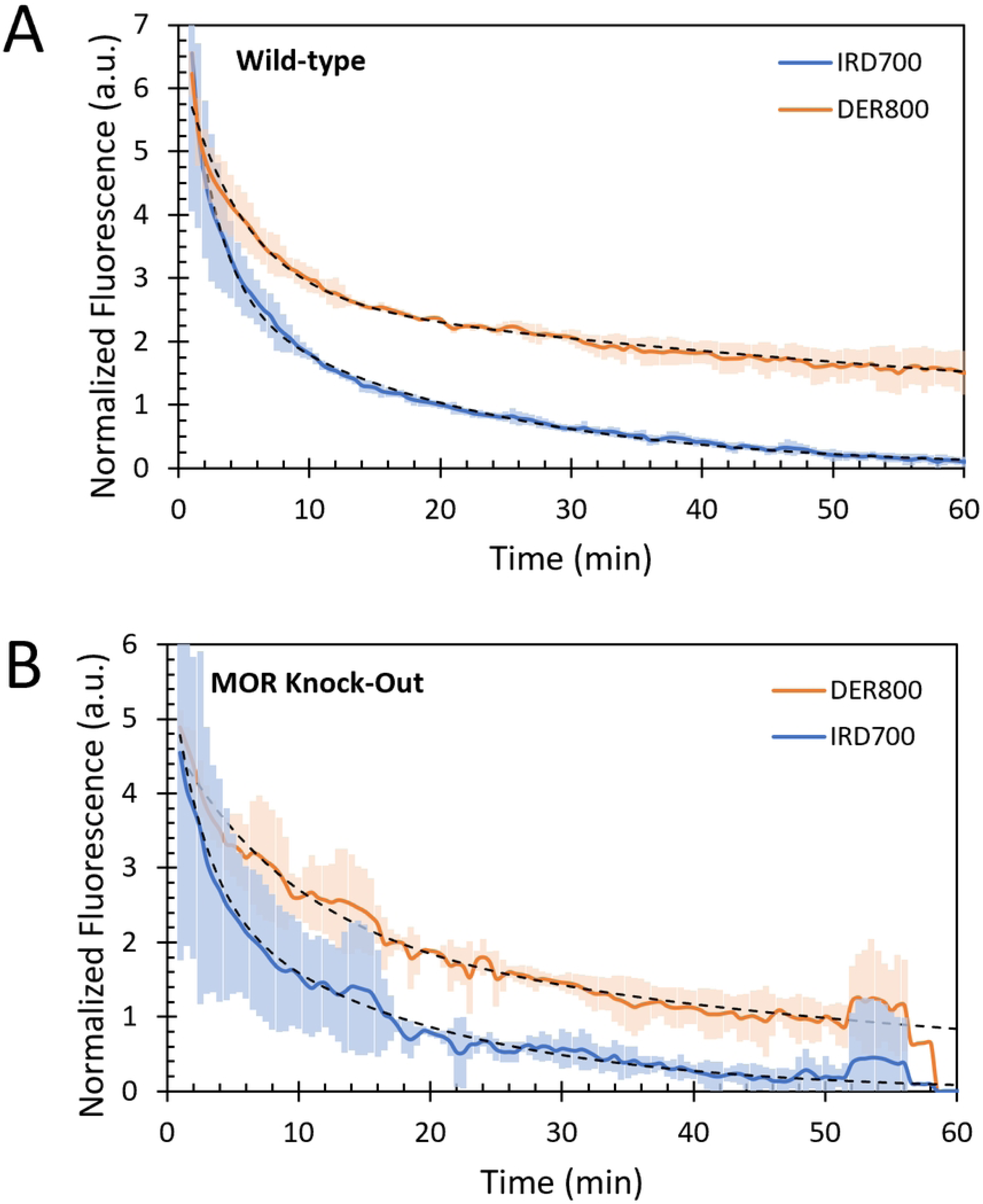
Plasma curves. The arterial plasma concentration curves following co-injection of IRDye700DX and DRM-800 acquired in (A) wildtype (n = 4) and (B) MOR knockout (n = 4) mice. Dashed lines represent biexponential curves fit to the data.

### Whole body distribution of DRM-800 differs in wildtype and MOR-knockout mice

Cryo-macrotome images were acquired from mice sacrificed 1 hour following injection of DRM-800 and IRDye^®^700DX in the manner described above. Full-resolution images were loaded into ImageJ as a stack and RGB white-light images were used to locate the structures of interest. Figure 5 shows representative examples of fluorescent and RGB slices in a wildtype and knockout mice at levels corresponding to the spiral ganglia.

**Fig 5.**
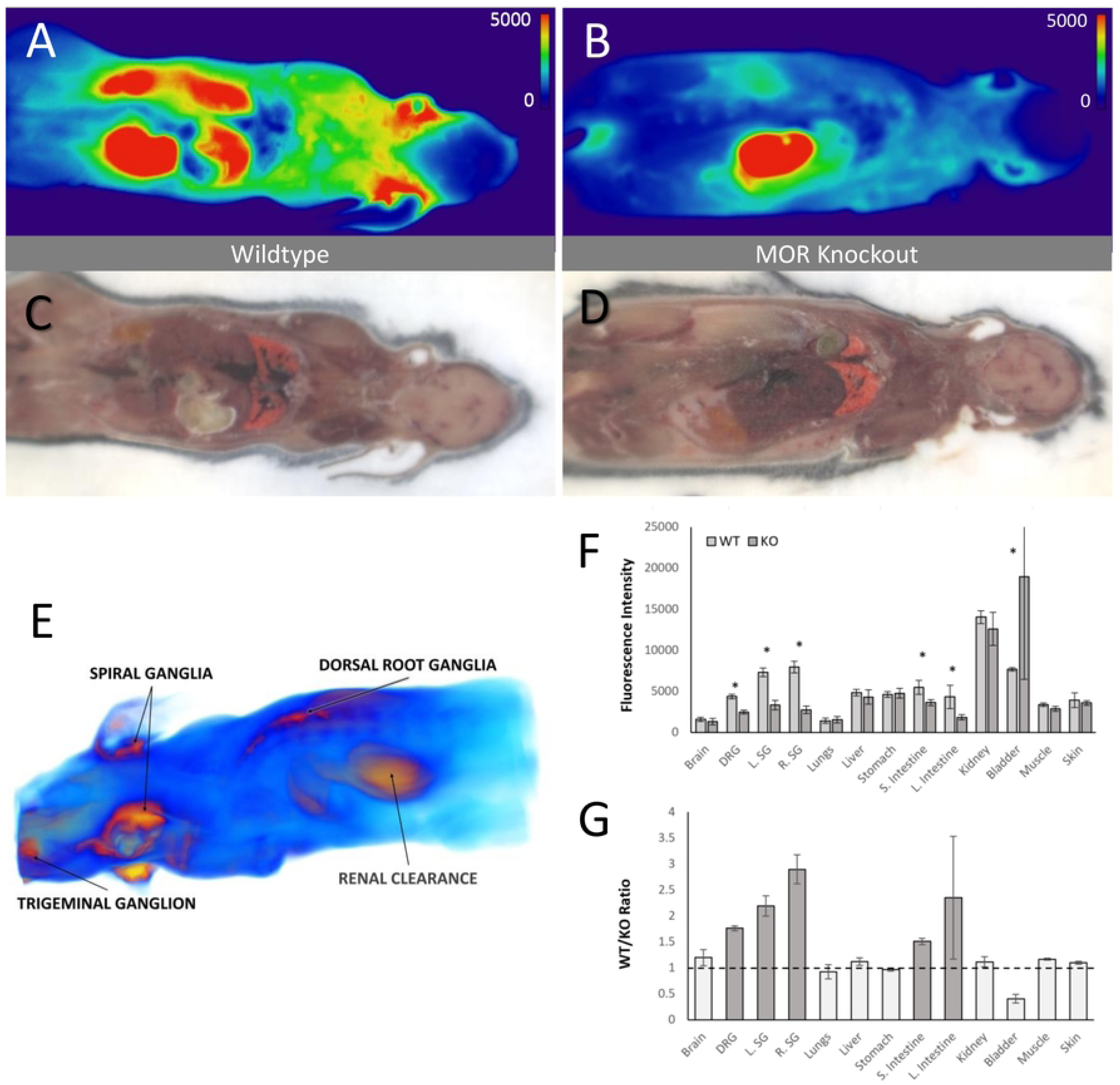
Biodistribution & Specific Binding. The DRM-800 fluorescence intensity (RFUs) in (A) wildtype and (B) MOR knockout mice 1 hour following injection, imaged on the cyro-macrotome, and the corresponding white-light color images (C-D) of the same slices showing the anatomical structures. The two mice were imaged under the same conditions and calibrated so that comparisons can be made between animals. (E) Fluorescence intensity data acquired from each slice was thresholded, stacked and volume-rendered using ParaView. (F) The fluorescence intensity in wildtype (WT; light grey bars) and MOR knockout (KO; dark grey bars) mice at thirteen different structures of interest. Asterisks indicate significant difference (p < 0.05). (G) The ratio of fluorescence between these two groups at the structures of interest, with solid grey bars indicating regions with a ratio significantly higher than unity. (DRG - dorsal root ganglia; SG - spiral ganglion; TG - trigeminal ganglion).

Figure 5E shows a volume rendering of the fluorescence signal, visualizing the highest signal regions. Significantly, regions that were well above the median level of fluorescence and that qualitatively could be perceived by adjusting the opacity and colormap thresholds were regions corresponding to the spiral ganglia, dorsal root ganglia, and trigeminal ganglion located at the skull base, as well as clearance by the kidneys.

Region-of-interest analysis was performed on slices such as those in Figure 5A-B, to determine the average fluorescence intensity in wildtype and knockout mice, as well as the ratio of intensity between the wildtype and knockout mice averages. A ratio greater than unity corresponds to specific DRM-800 binding to MOR, since the animals differ only in the presence or absence of MOR and represents an important test for whether DRM-800 is an effective biomarker for the presence or absence of available MOR. Figure 5F summarizes the fluorescence intensity for 13 different structures: brain, dorsal root ganglion, left spiral ganglion, right spiral ganglion, trigeminal ganglion, lungs, liver, stomach, small and large intestine, kidney, muscle and skin. Asterisks indicate pairs of values that differ significantly (*p* < 0.05).

Figure 5G presents the ratio of average fluorescence values determined from the 13 structures of interest between wildtype and knockout mice. The highest difference between wildtype and knockout groups was observed in the trigeminal ganglion (3.1 ± 0.2) and spiral ganglia (2.9 ± 0.3 and 2.2 ± 0.2, for right and left, respectively) followed by the dorsal root ganglion (1.8 ± 0.05). The small and large intestine regions of wildtype mice also showed elevated levels of DRM-800 compared with knockout mice.

### In Silico Specific Uptake and Internalization

As a corollary to the *in vitro* and *in vivo* animal studies presented in this paper, the Authors present numerical experiments that demonstrate the potential of DRM-800 to be used as a *dynamic multi-agent fluorescence (DMF)* biomarker. This represents one particular application of DRM-800 given the high affinity and high internalization rate observed in these studies. A three-tissue compartment model was used to describe the time-dependent distribution of DRM-800 in the vascular, extravascular free, bound and intracellular spaces. Figure 6 summarizes the compartment model used in the numerical simulations, which were performed for theoretical tissue structure of interest of a volume of 123 mm^3^ and containing about 42,000 ganglion neurons, which is consistent with a structure like the spiral ganglion. rate constants used in the simulation were *K*_1_ = 0.3 min^−1^, *k*_2_ = 0.01 min^−1^, *k*_4_ = 0.04 min^−1^, and *k*_5_ = 0.01 min^−1^, and *k*_3_ = BP_ND_ x *k*_4_, Binding potential was estimated, based on the *B*_max_ and *K*_d_ inferred from *in vitro* assays, to be approximately 10. The relationship between buprenorphine dose and binding potential (Fig. 6B) was modeled after reported MOR occupancy rates measured by PET imaging (18). Figure 6C shows the tissue fluorescence curves for DRM-800 and, for purposes of comparison, two other fluorophores: fluorescein-labeled naloxone (NLX-500) that binds but does not internalize, and IRDye^®^700DX (IRD-700) that doesn’t bind. The binding potential of DRM-800 in this case is 9.0. Figure 6D shows simulated DRM-800 curves for MOR availabilities from 6% to 90%, showing the relationship between the uptake slope and binding potential (or MOR availability).

**Fig. 6.**
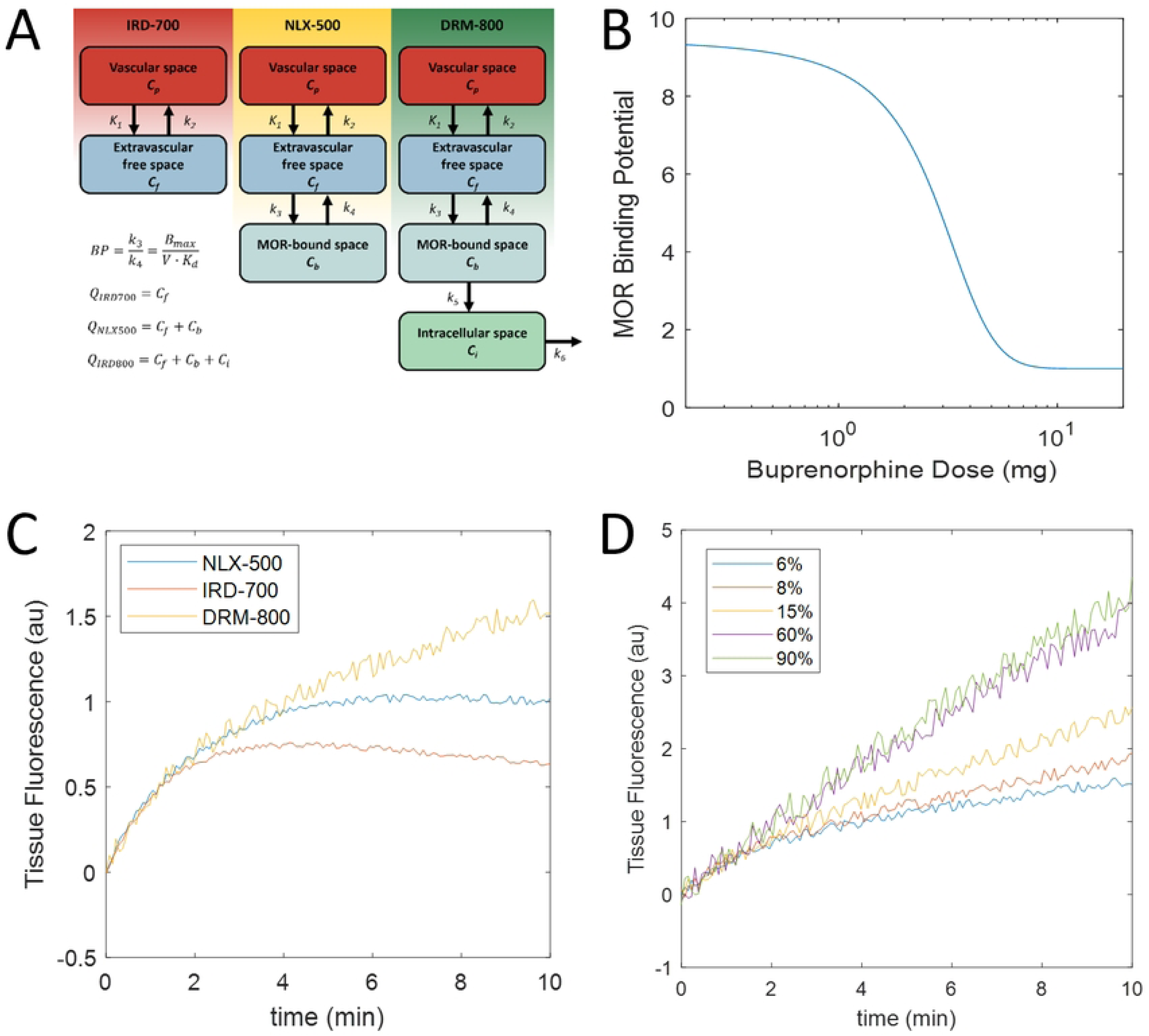
Tracer kinetic modeling. (A) Compartment models can describe the behaviour of IRDye^®^700DX (IRD-700), naloxone-fluorescein (NLX-500), and DRM-800. (B) Relationship between buprenorphine dose and MOR binding potential used in forward simulations (C) Tissue fluorescence as a function of time for the three agents (D) The DRM-800 fluorescence signal for different MOR occupancies.

## Discussion

In this paper, we describe a new biomarker of mu-opioid receptor occupancy that could enable individualized pharmacotherapy in patients with opioid use disorder. The molecule developed in this study, DRM-800, is a synthetic peptide conjugated to a GMP-available dye, IRDye®800CW. In contrast to previously developed fluorescent-labeled opioid peptides (23,24), DRM-800 is a high-affinity agonist that is rapidly internalized and, importantly, because it is developed using a GMP-produced and clinically dye, it can be moved along a pipeline that would enable exploratory investigational new drug (eIND) approval by the Food and Drug Administration (FDA) and first-in-human Phase 0 trials. However, before further progress along this pipeline is justified, DRM-800’s bioavailability, stability, binding efficacy and *in vivo* uptake must be established.

*In vitro* analysis of competitive binding demonstrates that DRM-800 is both an efficacious ligand to MOR and is rapidly internalized within minutes of binding to MOR. In particular, when MOR+ CHO-K1 cells were incubated with [Lys7]Dermorphin and DRM-800, the IC50 obtained for a range of DRM-800 concentrations indicated that about a 20x higher dose of [Lys7]Dermorphin is required to displace 50% of bound DRM-800. This was somewhat surprising because [Lys7]Dermorphin is one of the highest-affinity selective MOR ligands known. Furthermore, it isn’t entirely clear why the addition of IRDye^®^800CW would increase the affinity of the peptide. It is also possible that the internalization plays a role by reducing the molecules available for competition, and effectively reducing the *k_off_*rate. Naloxone displaced 50% of bound DRM-800 at a concentration 14x lower when the drugs were incubated simultaneously, however, when naloxone was added 20 minutes after DRM-800, its ability to competitively inhibit DRM-800 binding was completely eliminated. Both of these characteristics contribute to the potential for DRM-800 as a biosensor. In particular, the effect of internalization was observed by sequentially adding naloxone, which had no effect on specific binding signal at any concentration, since DRM-800 was already internalized and inaccessible. It was further corroborated by *in vitro* fluorescence microscopy images obtained for cells incubated at 35C with no sucrose, and 4C with 0.35 M sucrose, the latter condition being known to prevent internalization (21,22). As expected, internalization was readily observed by the presence of puncta at 35C, but not seen appreciably in the cells incubated with sucrose at 4C. This is in contrast to previously reported dermorphin-derived fluorescent constructs, which did not observe internalization behaviour when AlexaFluor 488, a dye with similar size and charge properties, was used (23). Given the complexity of the relationship between dye selection and subsequent agonist/internalization behaviour, the apparent rapid internalization of DRM-800 was serendipitous.

*In vivo* analysis of plasma concentration following intravenous injection demonstrates that DRM-800 exhibits small-molecule plasma kinetics behaviour similar to observed behaviour in other dye-small peptide conjugates (10). This is a beneficial property for a biomarker, especially if coupled with high receptor affinity and internalization, because the background signal (non-specific signal) will clear rapidly in the minutes to hours following injection, enhancing the MOR-specific contrast. It is interesting to note the differences in distribution behaviour of DRM-800 in wildtype vs. knockout mice; despite the large volume difference between Cp and Cf+Cb, because of the ubiquity of MOR across the body, it is possible that binding could enhance the egress of DRM-800 from the intravascular space, resulting in a higher distribution rate constant, *b*. Whole body distribution differences confirmed one of the main hypotheses we developed prior to this validation study: that opioid-rich ganglion would have sufficient expression of MORs to exhibit appreciable differences in fluorescence signal compared to knockout mice lacking any MOR expression. This was confirmed by leveraging a new imaging cryo-macrotome developed and maintained at Dartmouth College, which allows whole-body distribution to be imaged volumetrically by sequentially slicing through a frozen animal in 150 µm-thick sections. In addition to volumetric renderings enabling quick qualitative identification of structures and regions with signal enhancement, quantification of fluorescence (including up to four unmixed fluorophores and characterization of autofluorescence) is achievable by ROI analysis on acquired slices.

Our biosensing strategy, in order to avoid the need to cross the blood-brain barrier, is to target structures outside the CNS, and therefore we focused on peripheral structures known to be rich in MOR expression. Elevated MOR-specific signal was seen in the three ganglia regions of interest, as well as in the intestine, a region of known MOR expression, evinced by the gastrointestinal side effects attributed to opioid use. While the utility of the trigeminal ganglion, which transmits sensory afferents from the whiskers, is rodent-specific, the spiral ganglia and dorsal root ganglia are relevant in human disease. In particular, the identification of peripheral or accessible structures that may be useful as biomarker surrogates of MOR occupancy is a critical step in the development of a point-of-care tool.

As expected, we did not observe any appreciable signal in either groups of mice, corresponding to the brain. While IRDye^®^800CW is only 1.2 kDa in size, it is enough to prevent crossing the blood brain barrier. The unsuitability of DRM-800 to interrogate CNS structures implicated in OUD such as the amygdala and thalamus may be considered a limitation. However, the goal of this project is to develop a non-invasive point-of-care tool— medical imaging such as MRI and PET can already be used to interrogate internal brain structures non-invasively, but they are not appropriate for widespread, point-of-care use, due to their cost and complexity. Therefore, we sought a target that can serve as a surrogate for MOR occupancy and pharmacotherapeutic-induced changes in MOR dynamics and that is accessible *non-invasively*. The spiral ganglia are MOR-abundant bipolar neurons that innervate the organ of Corti, distally, and proximally project to the cochlear nuclei of the brainstem. Therefore, they are outside the blood brain barrier, and can be imaged otoscopically through the ear canal. To the best of our knowledge, this structure has not yet been exploited for biosensing and represents the innovative step of this project.

A critical limitation in this study is that the optimal time for imaging post-injection is unknown. Given the plasma concentration curves in Figure 4 we know that long imaging times on the order of 10-24 hrs are unlikely to result in adequate signal regardless of contrast between structures of interest and the background. However, this study does not present dynamic levels of fluorescence as a function of time after injection, due to the destructive nature of the cryo-macrotome process. In future experiments, we will focus on the structures that were elevated compared with knockout mice (e.g., DRG, SG) and measure the time-concentration curves of DRM-800 following injection to better understand the pharmacodynamics associated with DRM-800’s interaction with these sites. The reader may be skeptical of the idea that an external structure not related to higher-order neural processing might provide any helpful information to a disease characterized by complex psychosocial determinants. In fact, it is the highly individualized nature of OUD that beseeches an objective, pharmacological tool to characterize a patient-specific dose-response curve to pharmacotherapy. Subsequent animal experiments are underway to determine whether changes in peripheral MOR expression and internalization behavior coincide with initial exposure to morphine as well as during taper-down or rapid naloxone-induced withdrawal*. In silico* results presented in this paper demonstrate buprenorphine-mediated changes in DRM-800 fluorescence concentration curves under assumed parameters representative of in situ human ganglion measurements. These suggest at least the possibility that DRM-800 dynamic fluorescence could act as a surrogate for binding potential, and therefore by extension, occupancy of MOR by a pharmacotherapeutic agent like buprenorphine. Future studies are planned to further develop the feasibility of this concept.

## Conclusions

This study characterizing DRM-800 pharmacokinetics and evaluating the feasibility as a biomarker for MOR occupancy demonstrates its potential for further development as an *in vivo* assay. The pharmacokinetics and pharmacodistribution suggest specific uptake with structures known to express high levels of MOR. Competitive binding kinetics and rapid internalization suggest that detectable levels of fluorescence that correlate to MOR availability could be induced in those structures, and thus serve as a quantitative tool and surrogate for clinically relevant structures, with the goal of individualizing medical treatment of opioid use disorder.

## Acknowledgements

This work was funded by an NIH/NIBIB research grant (R21EB024771), and involved methodology developed by NIH/NCI grants R00CA190890 (JTE), R01CA184354 (SCD), and R01CA188491 (SCD). The corresponding author discloses intellectual property related to the work published in this manuscript in the form of US provisional patent application No. 62/933,644.

## Notes

### Competing Interest Statement

I have read the journal's policy and the authors of this manuscript have the following competing interests: The corresponding author discloses intellectual property related to the work published in this manuscript in the form of US provisional patent application No. 62/933,644.

